# Targeting DNA Damage Response and Replication Stress in Pancreatic Cancer

**DOI:** 10.1101/713545

**Authors:** Stephan B. Dreyer, Rosie Upstill-Goddard, Viola Paulus-Hock, Clara Paris, Eirini-Maria Lampraki, Eloise Dray, Bryan Serrels, Giuseppina Caligiuri, Selma Rebus, Holly Brunton, Richard Cunningham, Craig Nourse, Ulla-Maja Bailey, Marc Jones, Kim Moran-Jones, Derek W. Wright, Fraser Duthie, Karin Oien, Lisa Evers, Colin J. McKay, Grant A. McGregor, Aditi Gulati, Rachel Brough, Ilirjana Bajrami, Stephan Pettitt, Michele L. Dziubinski, Juliana Candido, Frances Balkwill, Simon T. Barry, Robert Grützmann, Lola Rahib, Glasgow Precision Oncology Laboratory, Australian Pancreatic Cancer Genome Initiative, Amber Johns, Marina Pajic, Fieke E. M. Froeling, Phillip Beer, Elizabeth A. Musgrove, Gloria M. Petersen, Alan Ashworth, Margaret C. Frame, Howard C. Crawford, Diane M. Simeone, Chris Lord, Debabrata Mukhopadhyay, Christian Pilarsky, Susanna L. Cooke, Nigel B. Jamieson, Jennifer P. Morton, Owen J. Sansom, Peter J. Bailey, Andrew V. Biankin, David K. Chang

**Affiliations:** Wolfson Wohl Cancer Research Centre, Institute of Cancer Sciences, University of Glasgow, Garscube Estate, Switchback Road, Bearsden, Glasgow, Scotland G61 1QH, UNITED KINGDOM; West of Scotland Pancreatic Unit, Glasgow Royal Infirmary, Glasgow G31 2ER UNITED KINGDOM; Cancer Research UK Beatson Institute, Garscube Estate, Switchback Road, Glasgow G61 1BD, UNITED KINGDOM; Clara Paris; UT Health San Antonio Department of Biochemistry and Structural Biology San Antonio, TX 78229-3900 U.S.A.; MRC Institute of Genetics & Molecular Medicine, Edinburgh Cancer Research UK Centre, University of Edinburgh, Crewe Road South, Edinburgh, EH4 2XR, UNITED KINGDOM; Stratified Medicine Scotland, Queen Elizabeth University Hospital, Glasgow, G51 4TF, UNITED KINGDOM; College of Medicine, Veterinary, and Life Sciences, University of Glasgow, Glasgow G12 8TA, UNITED KINGDOM; Department of Pathology, Queen Elizabeth University Hospital, Glasgow, G51 4TF, UNITED KINGDOM; Greater Glasgow and Clyde Bio-repository, Pathology Department, Queen Elizabeth University Hospital, 1345 Govan Road, Glasgow, G51 4TY, UNITED KINGDOM; CRUK Gene Function Laboratory and Breast Cancer Now Toby Robins Research Centre, The Institute of Cancer Research, Fulham Road, London, SW3 6JB, UNITED KINGDOM; Department of Molecular and Integrative Physiology, University of Michigan, 1137 E. Catherine St., An Arbor, MI, 48109, USA; Barts Cancer Institute, Queen Mary University of London, Charterhouse Square, London, EC1 M6BQ, UNITED KINGDOM; Bioscience, Oncology, IMED Biotech Unit, AstraZeneca, Cambridge, UNITED KINGDOM; Department of Surgery, Universitätsklinikum Erlangen, Erlangen, GERMANY; Pancreatic Cancer Action Network, Manhattan Beach, California, USA; The list of Glasgow Precision Oncology Laboratory and their affiliations is at the end of the paper; The list of Australian Pancreas Genome Initiative and their affiliations is at the end of the paper; The Kinghorn Cancer Centre, 370 Victoria Street, Darlinghurst and Garvan Institute of Medical Research, Sydney, NSW 2010, AUSTRALIA; Epigenetics Unit, Department of Surgery & Cancer, Imperial College London, Hammersmith Campus, Du Cane Road, London, W12 0NN, UNITED KINGDOM; Lustgarten Foundation Pancreatic Cancer Research Laboratory, Cold Spring Harbor Laboratory, Cold Spring Harbor, NY, USA, 11724; Sanger Institute, Wellcome Genome Campus, Cambridge, UNITED KINGDOM; Mayo Clinic, Rochester, Minnesota, 55905, USA; UCSF Helen Diller Family Comprehensive Cancer Center, San Francisco, California, 94158, USA; Pancreatic Cancer Center, Perlmutter Cancer Center, NYU Langone Health, New York, NY 10016, USA; Department of Biochemistry and Molecular Biology, Mayo Clinic College of Medicine and Science, Jacksonville, FL 32224, USA; South Western Sydney Clinical School, Faculty of Medicine, University of NSW, Liverpool NSW 2170, Australia; Glasgow Precision Oncology Laboratory, University of Glasgow, Institute of Cancer Sciences, Wolfson Wohl Cancer Research Centre, Garscube Estate, Switchback Road, Glasgow, G61 1QH, United Kingdom; West of Scotland Pancreatic Unit, Glasgow Royal Infirmary, Glasgow, G31 2ER, United Kingdom; Department of Pathology, Southern General Hospital, Greater Glasgow & Clyde NHS, Glasgow, G51 4TF, United Kingdom; West of Scotland Genetic Services, NHS Greater Glasgow and Clyde, Queen Elizabeth University Hospital Campus, Glasgow, G51 4TF, United Kingdom; The Kinghorn Cancer Centre, Garvan Institute of Medical Research, 370 Victoria Street, Darlinghurst, Sydney, New South Wales 2010, Australia; Queensland Centre for Medical Genomics, Institute for Molecular Bioscience, University of Queensland, St Lucia, Queensland 4072, Australia; Royal North Shore Hospital, Westbourne Street, St Leonards, New South Wales 2065, Australia; Bankstown Hospital, Eldridge Road, Bankstown, New South Wales 2200, Australia; Liverpool Hospital, Elizabeth Street, Liverpool, New South Wales 2170, Australia; Westmead Hospital, Hawkesbury and Darcy Roads, Westmead, New South Wales 2145, Australia; Royal Prince Alfred Hospital, Missenden Road, Camperdown, New South Wales 2050, Australia; Fremantle Hospital, Alma Street, Fremantle, Western Australia 6959, Australia; Sir Charles Gairdner Hospital, Hospital Avenue, Nedlands, Western Australia 6009, Australia; St John of God Healthcare, 12 Salvado Road, Subiaco, Western Australia 6008, Australia; Royal Adelaide Hospital, North Terrace, Adelaide, South Australia 5000, Australia; Flinders Medical Centre, Flinders Drive, Bedford Park, South Australia 5042, Australia; Greenslopes Private Hospital, Newdegate Street, Greenslopes, Queensland 4120, Australia; Envoi Pathology, 1/49 Butterfield Street, Herston, Queensland 4006, Australia; Princess Alexandria Hospital, Cornwall Street & Ipswich Road, Woolloongabba, Queensland 4102, Australia; Austin Hospital, 145 Studley Road, Heidelberg, Victoria 3084, Australia; Victorian Cancer Biobank, 1 Rathdown Street, Carlton, Victoria 3053, Australia; Johns Hopkins Medical Institute, 600 North Wolfe Street, Baltimore, Maryland 21287, USA; ARC-NET Center for Applied Research on Cancer, University of Verona, Via dell’Artigliere, 19 37129 Verona, Province of Verona, Italy; University of California, San Francisco, 500 Parnassus Avenue, San Francisco, California 94122, USA; Wolfson Wohl Cancer Research Centre, Institute of Cancer Sciences, University of Glasgow, Garscube Estate, Switchback Road, Bearsden, Glasgow, Scotland G61 1BD, United Kingdom; Greater Glasgow and Clyde National Health Service, 1053 Great Western Road, Glasgow G12 0YN, United Kingdom

## Abstract

Continuing recalcitrance to therapy cements pancreatic cancer (PC) as the most lethal malignancy, which is set to become the second leading cause of cancer death in our society. We interrogated the transcriptome, genome, proteome and functional characteristics of 61 novel PC patient-derived cell lines to define novel therapeutic strategies targeting the DNA damage response (DDR) and replication stress. We show that patient-derived cell lines faithfully recapitulate the epithelial component of pancreatic tumors including previously described molecular subtypes. Biomarkers of DDR deficiency, including a novel signature of homologous recombination deficiency, co-segregates with response to platinum and PARP inhibitor therapy *in vitro* and *in vivo*. We generated a novel signature of replication stress with potential clinical utility in predicting response to ATR and WEE1 inhibitor treatment. Replication stress and DDR deficiency are independent of each other, creating opportunities for therapy in DDR proficient PC, and post-platinum therapy.

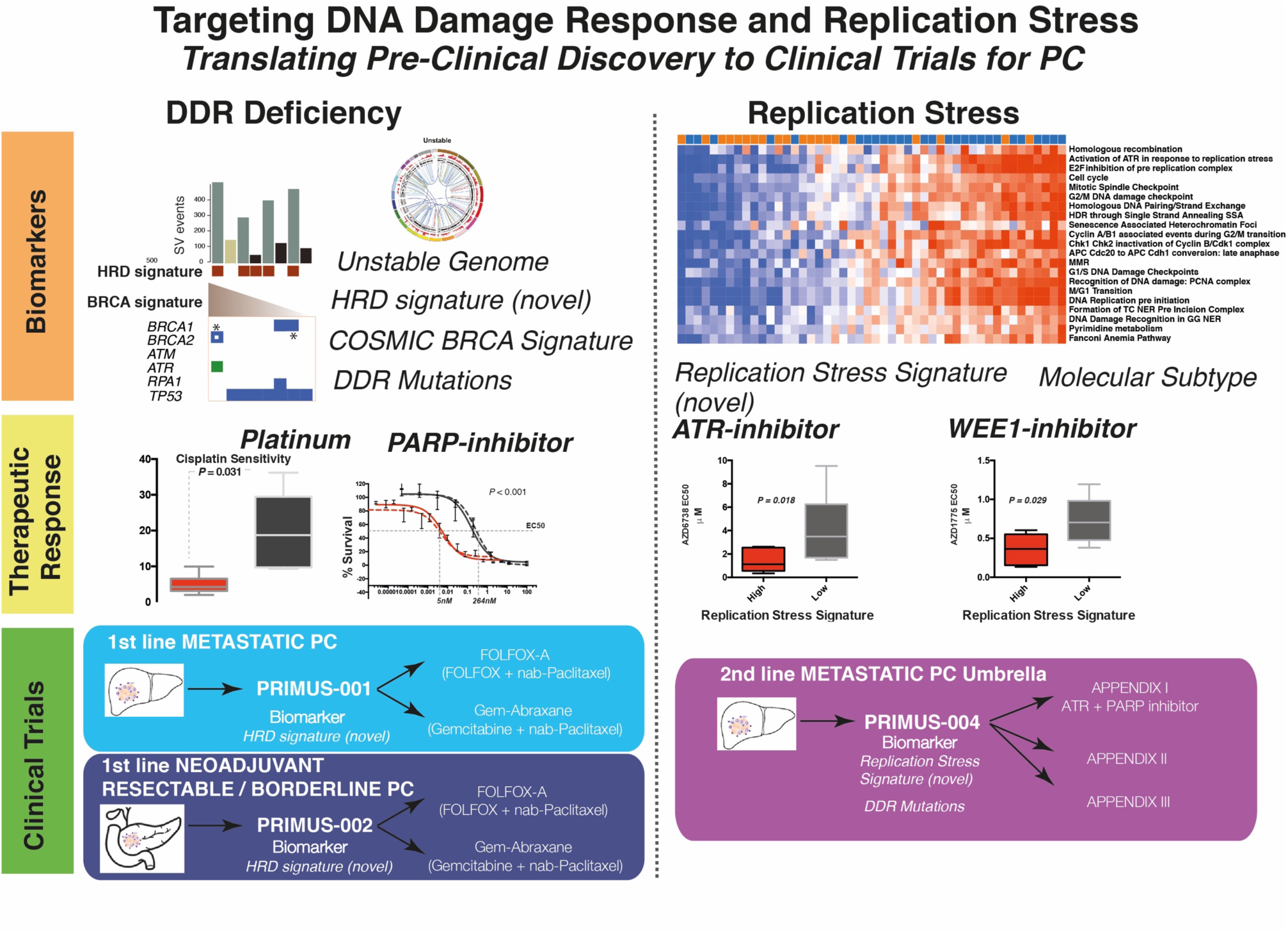

**STATEMENT OF SIGNIFICANCE:** We define therapeutic strategies that target subgroups of PC using novel signatures of DNA damage response deficiency and replication stress. This potentially offers patients with DNA repair defects therapeutic options outside standard of care platinum chemotherapy and is being tested in clinical trials on the Precision-Panc platform.

## INTRODUCTION

Pancreatic Cancer (PC) has recently overtaken breast cancer to become the third leading cause of cancer death in the USA^1^ and is predicted to become the second within a decade^2^. Pancreatic ductal adenocarcinoma (PDAC), the more common form of PC, is dominated by mutations in four well-known cancer genes (*KRAS, TP53, CDKN2A and SMAD4*). Only a few genes are mutated in 5 – 15% of cases, amidst an ocean of infrequently mutated genes in the majority of patients ^3–11^. This diversity may explain the lack of progress with targeted therapies, as actionable genomic events being targeted therapeutically are present in only a small proportion of unselected participants in clinical trials^12^. To better select patients to clinical trials, biomarkers that predict response to novel, and established treatments are urgently needed and must extend beyond the detection of point mutations in coding genes and low prevalence actionable genomic events.

Whilst molecular subtyping of cancer based on biological attributes can facilitate drug discovery, to be clinically relevant, the optimal taxonomy must inform patient management through prognostication or more importantly, treatment selection^13^. Recent studies have subtyped PC in various ways ^5,9,14–18^, grouping similarities based on structural attributes of genomes, genes mutated in pathways, or molecular mechanisms inferred through mRNA expression. Despite discrepancies in nomenclature, one molecular class (variably termed quasi-mesenchymal, basal-like or squamous) is consistently defined and is associated with a poor prognosis^19,20^. These biologically based molecular taxonomies of PC, whilst associated with differences in outcome, are yet to inform treatment decisions.

DNA damage response (DDR) deficiency is a hallmark of cancer, including PC^8^, which is thought to render some tumors preferentially sensitive to DNA damaging agents such as platinum. There is a growing compendium of novel therapeutics that target DNA damage response mechanisms and the cell cycle such as ATR and WEE1 inhibitors^21^. Genomic instability, a key feature of many cancers, typically secondary to defects in DNA replication and repair during the cell cycle often results in replication stress^22,23^. Oncogene activation drives replication stress, particularly through RAS and MYC signaling, both of which are prevalent molecular features of PC ^22,24,25^. The platinum containing regimen, FOLFIRINOX, has become the standard of care for all stages of PC, yet is only suitable for patients with good performance status, however the majority of patients unfortunately do not respond^26–28^. Consequently, many patients suffer the morbidity, and even mortality, of systemic platinum chemotherapy with little or no survival benefit or quality of life. Biomarker driven patient selection strategies, and novel therapeutics that build on platinum response or disease stabilization that target DDR mechanisms, provide a substantial opportunity to improve outcomes.

Here we use 61 patient-derived cell lines (PDCL) of PC (Supplementary Table 2), to define subtype-specific molecular mechanisms and identify opportunities for molecular subtype-directed treatment selection that targets DDR mechanisms. We performed mRNA expression analysis (RNAseq) (*n* = 48) complemented with whole genome sequencing (*n* = 47), which was further enhanced with Reverse Phase Protein Arrays (RPPA), functional screens using small interfering RNA (siRNA) and targeted functional analysis. We identify novel biomarkers of DDR deficiency and replication stress with potential clinical utility that associate with therapeutic sensitivity. We show that DDR deficiency exists independently of replication stress, the previously identified poor prognostic Squamous subtype is enriched for replication stress and transcriptomic readouts of replication stress confer sensitivity to therapeutics that target the cell cycle checkpoint machinery.

## RESULTS

### Patient-derived cell lines recapitulate PC subtypes

We recently defined four transcriptomic subtypes of PC^5,19^, with two distinct primary lineages, termed Classical Pancreatic (which can be further divided into Pancreatic Progenitor, Immunogenic and Aberrantly Differentiated Endocrine eXocrine subtypes) and Squamous. A key distinction is the epigenetic profile of the Squamous subtype, with chromatin modification and methylation orchestrating the loss of pancreatic endodermal transcriptional networks, and as a consequence, suppressing transcripts that designate a pancreatic identity^5^ (Figure 1a). Hierarchical clustering of RNAseq data from the 48 PDCLs recapitulated the two primary classes of PC (Supplementary Figure 1). Twenty-eight (58%) of the PDCLs were classified as Squamous, and 20 (42%) were Classical (Supplementary Table 5). The preservation of the 26 transcriptional networks (Gene Programs) we previously described in bulk PC^5^ was compared to PDCL derived gene programs (Supplementary Figure 1, Supplementary tables 13 and 14). In total, 17 of the 26 Gene programs were closely recapitulated in the PDCLs (Supplementary Figure 1), with the expected absence of immune infiltrate related transcriptional networks (Supplementary Tables 14). The lack of stroma permitted higher resolution of epithelial transcriptomic networks, revealing key mechanisms that are difficult to discern form biopsy samples. Differential expression of genes related to DNA damage response, cell cycle control and morphogenic processes were observed between subtypes and correlated in both PDCLs and bulk tumor samples (Figure 1a). These findings suggest that PDCLs are representative of bulk PC and can be used to develop novel therapeutic strategies for the clinic, and that epithelial cell purity can provide greater sensitivity in detecting aberrant mechanisms.

**Figure 1.**
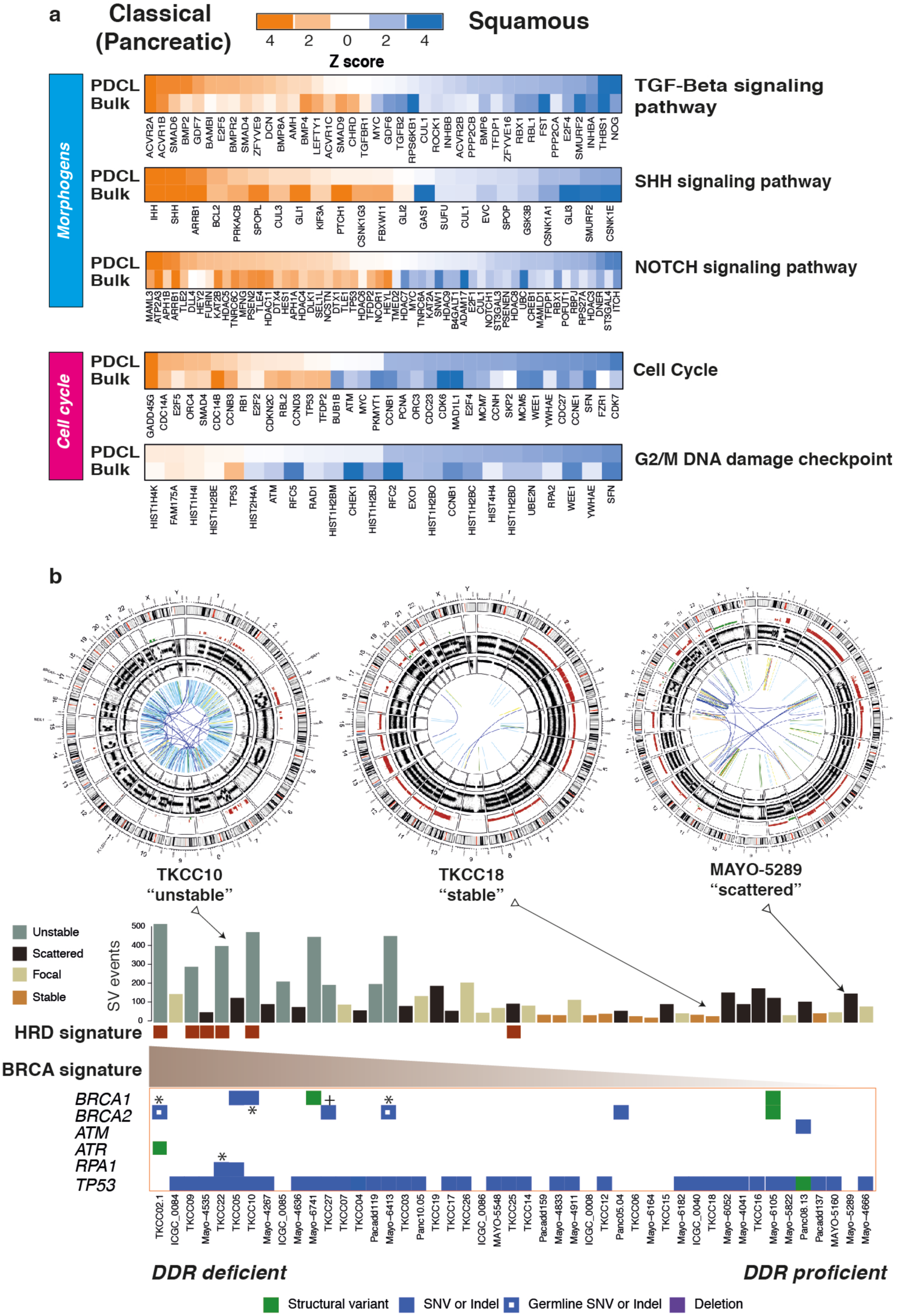
Subtype specific differences and DNA damage response in PDCLs of PC. **a)** Heatmaps of key genes in pathways important in carcinogenesis, grouped into distinct molecular processes related to morphogenesis and cell cycle control between molecular subtypes of bulk tumor and PDCLs of PC. The degree of color saturation is proportional to the degree of enrichment in the Squamous (blue) and Classical Pancreatic (orange) subtypes for all samples within each subtype, genes are ranked by most differentially expressed between subtypes. **b)** Surrogate biomarkers of DDR deficiency, defined by large scale sequencing projects of PC, include (1) unstable genome (> 200 SVs), (2) the novel homologous recombination deficiency (HRD) signature, (3) high ranking BRCA mutational signature, and (4) deleterious mutations in DDR pathway genes. PDCLs are ranked from left to right based upon the COSMIC BRCA mutational signature, with SV subtype, number of structural variations and HRD signature status symbolized on the top bar. Examples of Circos plots for 3 PDCLs are included, representing “unstable”, “stable” and “scattered” subtypes.

### Defining DDR deficiency in PDCLs of Pancreatic Cancer

Various biomarkers of DDR deficiency associated with therapeutic response have been proposed, but not validated and not used clinically in PC. Genomic markers of DDR deficiency such as a high ranking COSMIC BRCA point mutational signature co-segregates with high prevalence of structural variants, termed the ‘unstable’ genomic subtype, and deleterious mutations in homologous recombination repair (HR) pathway genes such as *BRCA1* and *2* and *PALB2* ^4^ (Figure 1b). We previously demonstrated that these signatures associate with clinical response to platinum in PC^4^. More recently, early data also suggest a therapeutic signal using PARP inhibitors, however, efficacy is not well defined beyond *BRCA1 / 2* mutations ^4,29,30^. To address this, we defined PDCLs as DDR deficient based on the interaction of 4 putative biomarkers: (1) structural variation number and pattern (> 200 SVs = unstable genome), (2) a high COSMIC BRCA mutational signature (ranked within top quintile), (3) a positive homologous recombination deficiency (HRD) signature^31,32^ and (4) mutations in key DDR genes (*BRCA1, BRCA2, ATM, ATR, RPA1, RAD51, RAD54, FANCA*) (FIGURE 1b, Supplementary Figure 2, Supplementary Table 10). Out of 47 PDCLs with WGS data, 6 (13%) had a positive HRD signature, 9 (19%) had > 200 SVs (unstable genome), 10 (21%) had mutations in DDR genes of which 2 were germline variants (both in *BRCA2*) (Figure 1b, Supplementary Table 11). There were 4 PDCLs with homozygous mutations in either *BRCA1, BRCA2* or *RPA1*, and these were all associated with unstable genomes, and 3 of these were HRD signature positive (Figure 1b, Supplementary Table 3). Significant overlap existed amongst these, with *n* = 3 PDCLs with all 4 biomarkers present, *n* = 1 had 3 (unstable genome, HRD signature, BRCA mutational signature) biomarkers positive, *n* = 4 were positive for 2 biomarkers and the remaining *n* = 8 had 1 biomarker positive (Supplementary Table 3).

**Figure 2.**
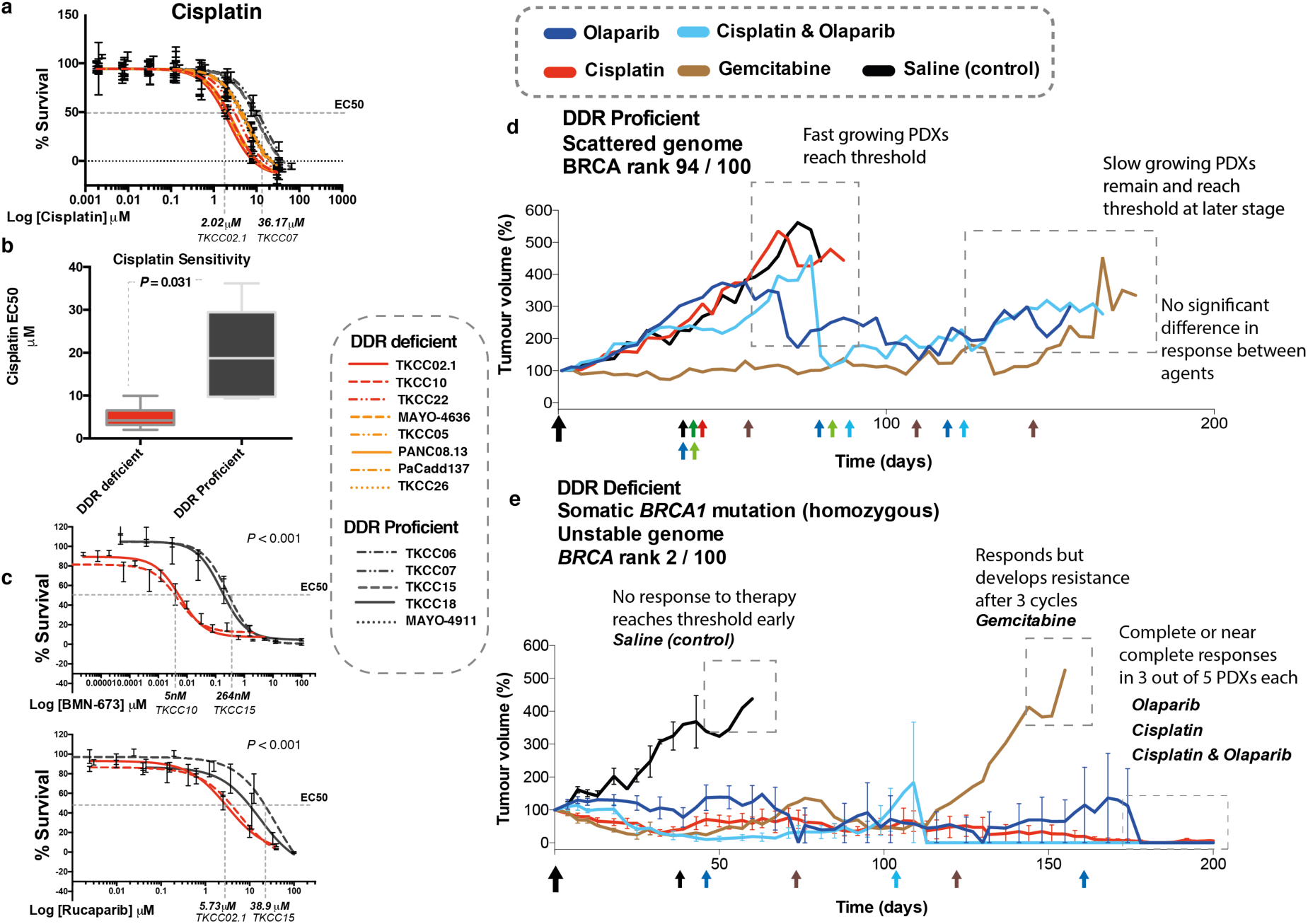
Targeting DDR deficient PC with Platinum and PARP inhibitors. **a**) Cell viability after 72 hours of Cisplatin treatment in PDCLs. Dotted line indicates EC50 in most sensitive PDCL was around 15 times more sensitive than most resistant PDCLs. **b**) Boxplot of mean Cisplatin EC50 in PDCLS stratified by DDR status. Box represents 95% confidence interval, and whiskers minimum and maximum range. *P* calculated using Mann Whitney test between mean EC50 in each group. **c)** PARP inhibitor (BMN-637 and Rucaparib) response in PDCLs. Dotted lines indicate EC50 between most sensitive and most resistant PDCLs. *P* indicates statistical difference between TKCC 10 (HRD signature positive) and TKCC15 (DDR proficient) using non-linear regression analysis. **d**) PDX 2133 and **e**) PDX 2179 (DDR deficient) treated with a panel of DNA damaging agents and Gemcitabine.

### DDR deficiency co-segregates with response to Platinum and PARP inhibitor treatment

To investigate the relationship between these putative biomarkers of DDR deficiency and Platinum and PARP inhibitor response, cell viability assays were performed on 15 PDCLs. PDCLs defined as DDR deficient were more sensitive to both Cisplatin therapy (*P* = 0.031) and PARP inhibition (*P* < 0.001) compared to DDR proficient PDCLs (Figure 2, Supplementary Table 11). The DDR deficient PDCLs all had EC50s to platinum of below the sensitivity threshold (10**μ**M) set by large scale pan-cancer cell line drug screens (*n* = 880) using Cisplatin (cancerrxgene.org (COSMIC)). These results suggest that DDR deficiency, as defined by these putative biomarkers have potential clinical utility in predicting response to platinum treatment. Importantly, this included both somatic and germline mutations, suggesting that therapeutic sensitivity extends beyond germline *BRCA1* and *2* mutations (Figure 2, Supplementary table 11).

To further define clinically applicable therapeutic response biomarkers of DDR deficiency *in vivo*, bulk tumor PDX models that represent both DDR proficient (PDX 2133) and deficient (PDX 2179) PC were generated in balb/c nude mice. The DDR proficient PDX did not respond to DNA damaging agents, including Cisplatin and the PARP inhibitor Olaparib combination (Figure 2). The DDR deficient PDX model, with a bi-allelic somatic *BRCA1* mutation, responded exceptionally to Cisplatin and Olaparib as monotherapy, and in combination (Figure 2), suggesting PARP inhibition can be as effective as Platinum chemotherapy in DDR deficient PC.

### Replication Stress is a feature of the Squamous subtype of PC

Replication Stress has been described to be closely related to DDR deficiency. As a consequence, we investigated subtype specific targeting of replication stress as a novel therapeutic strategy. We found significant subtype differences in the expression of genes controlling cell cycle, including the G2/M checkpoint in both PDCLs and bulk tumor PC (Figure 1). Expression of *WEE1* (*P* = 0.006), *CDK6* (*P* = 0.02) and *CDK7* (*P* < 0.001) was enriched in the squamous subtype in both PDCLs and bulk tumor (Figure 1a). We then used a combination of DNA maintenance, replication and cell cycle regulation network related transcriptional profiles from Gene Ontology (GO) and pathway enrichment analysis to define replication stress using mechanisms associated with DNA replication (ATR activation, chromosomal maintenance, E2F transcriptional pathways, homologous recombination, Fanconi anemia, base-excision repair, p53 signaling, ER stress, and RNA processing). This resolved into a transcriptomic signature (termed the Replication Stress signature) which was applied as a hypothetical biomarker of replication stress (Figure 3, Supplementary Table 12).

**Figure 3.**
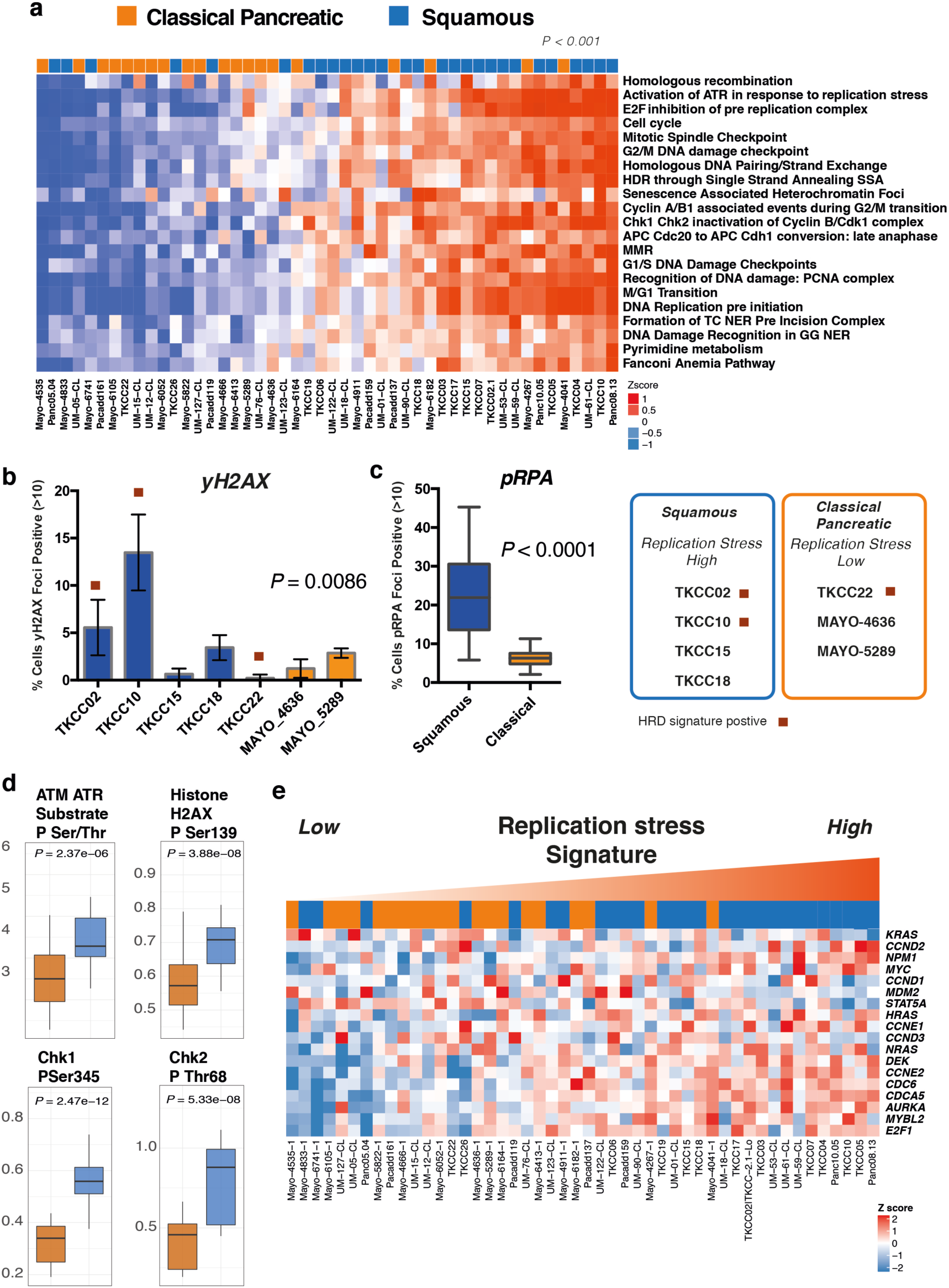
Replication Stress in PDCLs of PC. **a)** Heatmap of pathways and molecular processes (GO terms) involved in DNA maintenance and cell cycle regulation activated in replication stress and DNA damage response. PDCLs are ranked from right to left based on the decreasing novel transcriptomic signature score of replication stress and molecular subtype is indicated in the top bar demonstrating the association between activation of replication stress and the squamous subtype (*P* < 0.001, Chi square test, Low vs High). **b)** Immunofluorescent quantification of γH2AX and (**c**) pRPA at normal conditions, are elevated in the Squamous (blue), but not the Classical Pancreatic (orange) PDCLs. **d**) Proteomic analysis using RPPA of a panel of PDCLs demonstrated that replication stress response proteins are differentially activated in the squamous subtype. **e)** Heatmap demonstrating oncogene expression in PDCLs ranked from right to left by replication stress signature. Squamous PDCLs are enriched for oncogene activation and replication stress.

PDCLs with high replication stress were more likely to be of squamous subtype (*P* < 0.001) (Figure 3) and had significantly higher levels of pRPA at rest (a surrogate marker of single-stranded DNA break accumulation which infers replication stress) (*P* < 0.0001) (Figure 3). PDCLs with high replication stress and concurrent HR deficiency had a greater proportion of γH2AX positive cells at rest (a marker of double-stranded DNA breaks) (*P* = 0.0086) and had persistently high levels of pRPA and γH2AX positive cells at 20 hours after ionizing radiation, when pRPA and γH2AX should have returned to normal levels in cells with no replication defects and competent at repairing this level of DNA damage (Supplementary Figure 3). Reverse phase protein arrays (RPPA) inferred functional consequences with differential phosphorylation and activation of key effectors of DDR and cell cycle progression between Classical and Squamous subtypes (CHK1, CHK2, Rb, p21^CIP1/WAF1^, ATM/ATR substrates, cyclin D1, histone H2AX) (Figure 3, Supplementary Figure 3). The squamous subtype is also enriched for the activation and transcription of oncogenes including MYC and CCNE (Figure 3). Oncogene activation is known to cause replication stress secondary to genomic instability leading to activation of cell cycle checkpoint regulatory proteins involved in replication stress response such as ATR, WEE1 and CHK1^25,33^ (Supplementary Figure 4). These data demonstrate the squamous subtype is enriched for genes associated with replication stress, potentially secondary to oncogene activation, and provides a potential novel therapeutic strategy.

**Figure 4.**
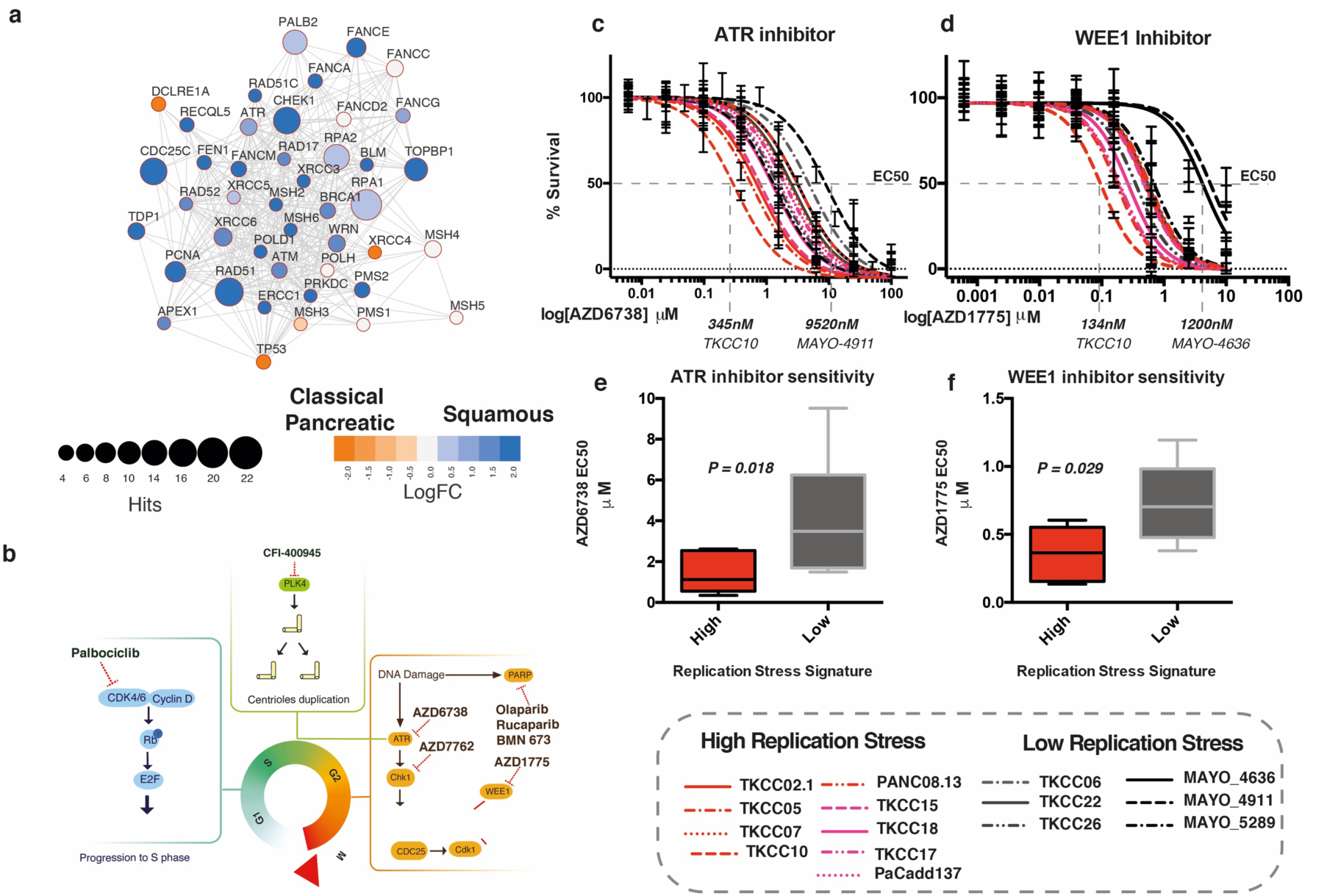
Targeting Replication Stress in PC. **a)** siRNA screen demonstrating transcriptome functional interaction (FI) sub-network, demonstrating preferential dependencies of cell cycle control and DNA maintenance genes in the squamous subtype. Different node colors represent dependencies in different molecular subtypes, and the size of each node is relative to the number of siRNA hits. **b)** Agents currently in clinical trial or approved for use in other cancer types that target cell cycle checkpoints. **c)** Dose response curves (EC50 shift) for ATR and **d)** WEE1 inhibitors calculated using MTS assay after 72 hours of drug treatment. **e)** and **f)** Mean relative EC50 for PDCLs stratified by replication stress score. Each boxplot represents mean EC50, box and whiskers represent minimum and maximum EC50 with 95% confidence interval. *P* calculated using Mann Whitney test between mean EC50 in each group

### Replication Stress is associated with sensitivity to Cell Cycle Checkpoint inhibitors

Differential expression of genes regulating the G2/M checkpoint in PDCLs and bulk tumors (such as *WEE1* and *CHEK1*) and the dependence on ATR activation in response to replication stress (Figure 3a) suggest that selective inhibition of these mechanisms may confer efficacy in tumors with high replication stress.

Further supporting this hypothesis, we performed an siRNA screen targeting genes controlling DNA damage repair and replication demonstrated a functional dependency on DNA damage response proteins, including ATM, ATR and CHK1 in squamous PDCLs (FIGURE 4a, Supplementary Figure 2). This is in keeping with the results from the immunofluorescent and RPPA analyses suggesting higher baseline levels of proteins associated with replication stress in the squamous PDCLs and a subsequent dependency on these proteins and cell cycle checkpoints for maintaining genomic integrity and cell survival.

Novel agents currently in early phase clinical trial targeting the cell cycle was used to generate a therapeutic testing strategy (Figure 4b) ^34–38^. Based on these data, in-vitro sensitivity was assessed using cell viability assays after a selection of PDCLs were treated with increasing doses of inhibitors of CHK1 (AZD7762), CDK4/6 (Palbociclib) and PLK4 (CFI-400945) demonstrating differential sensitivity (Supplementary Figure 4). Based on promising early clinical trial results in other cancer types ^37–42^, more extensive testing using inhibitors of ATR (AZD6738) and WEE1 (AZD1775) was performed on 15 PDCLs defined as high and low replication stress based on the replication stress signature score (Figure 3a, Supplementary Figure 4). This demonstrated that PDCLs with high replication stress were more sensitive to both ATR and WEE1 inhibition (Figure 4c-f). Importantly, these responses were independent of DDR status (Supplementary Figure 4), suggesting replication stress status is a more reliable marker of therapeutic response to ATR and WEE1 inhibition.

### Replication Stress is independent of DDR deficiency in PC

To further investigate the relationship of replication stress and DDR deficiency and the alignment of these therapeutic segments, a comparison was performed using the PDCL cohort. Using the described biomarkers of DDR deficiency and Replication Stress, a two by two grid was constructed to compare replication stress ranking and DDR deficiency (Figure 5). This demonstrated that signatures of DDR deficiency and replication stress are largely independent of each other, yet high replication stress is enriched in the squamous subtype (*P* = 0.007) (Figure 5). Therapeutic response data was overlapped based upon previously described experiments using ATR / WEE1 inhibitors and platinum, generating biomarker hypotheses for therapeutic responsiveness (Figure 5). PDCLs that are DDR deficient with high replication stress respond to both DDR targeting agents (e.g. Platinum and PARP inhibitors) and cell cycle checkpoint inhibitors (e.g. ATR and WEE1 inhibitors); DDR deficient, low replication stress to DDR agents only; DDR proficient, high replication stress to cell cycle checkpoint inhibitors only; and DDR proficient, low replication stress to neither class of agent (Figure 5).

**Figure 5.**
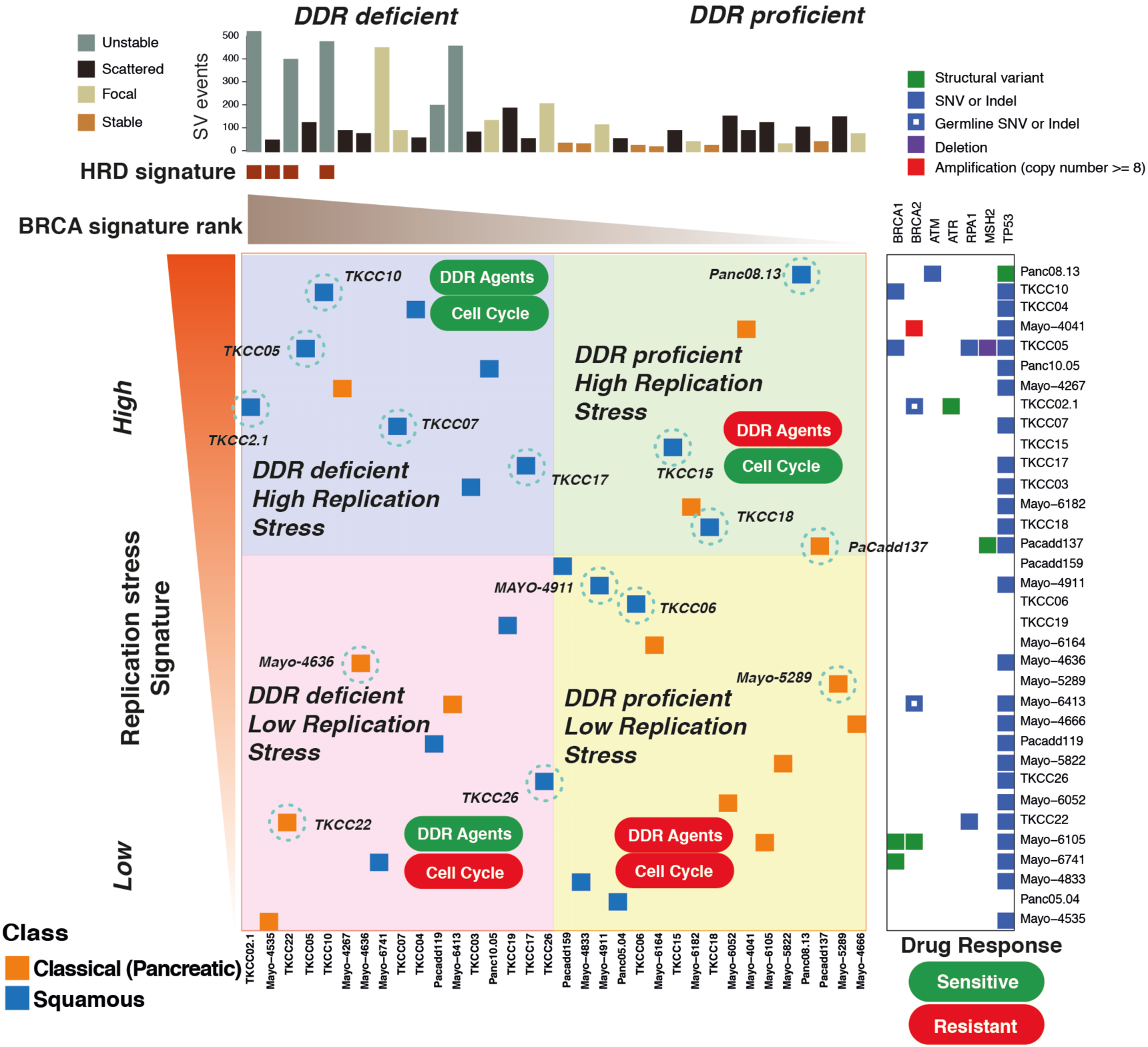
Relationship between DDR deficiency, Replication Stress and Therapeutic response in PDCLs of PC. PDCLs are ranked based on a novel transcriptomic signature of replication stress (y-axis), and a composite genomic readout of DDR deficiency (x-axis). DDR deficiency is a hierarchical score which incorporates the COSMIC BRCA mutational signature (signature 3), the number of structural variants distributed across the genome, and the HRD signature associated with BRCA deficiency. The combination of high/low states of each characteristic result in 4 groups. Squamous subtype PDCLs (blue squares) are associated with high replication stress (*P* = 0.007, Chi-squared). PDCLs tested are identified and encircled blue. DDR deficiency, and the replication stress signature predicted differential therapeutic response.

### Potential Clinical utility of the Replication Stress signature

To assess the potential clinical validity and utility of these pre-clinical data, the relationship between the replication stress signature score and molecular subtypes in bulk tumor samples was assessed using published transcriptomic data sets of PC^5^,^9^. This included whole transcriptome sequencing sets acquired through the International Cancer Genome Consortium (ICGC), totaling 94 patients with primary resected PC (Figure 6). This recapitulated the association between squamous molecular subtype and high replication stress (*P* = 0.006) (Figure 6, Supplementary Figure 5), with 50% of squamous tumors in the top quartile of tumors ranked by the replication stress score.

**Figure 6.**
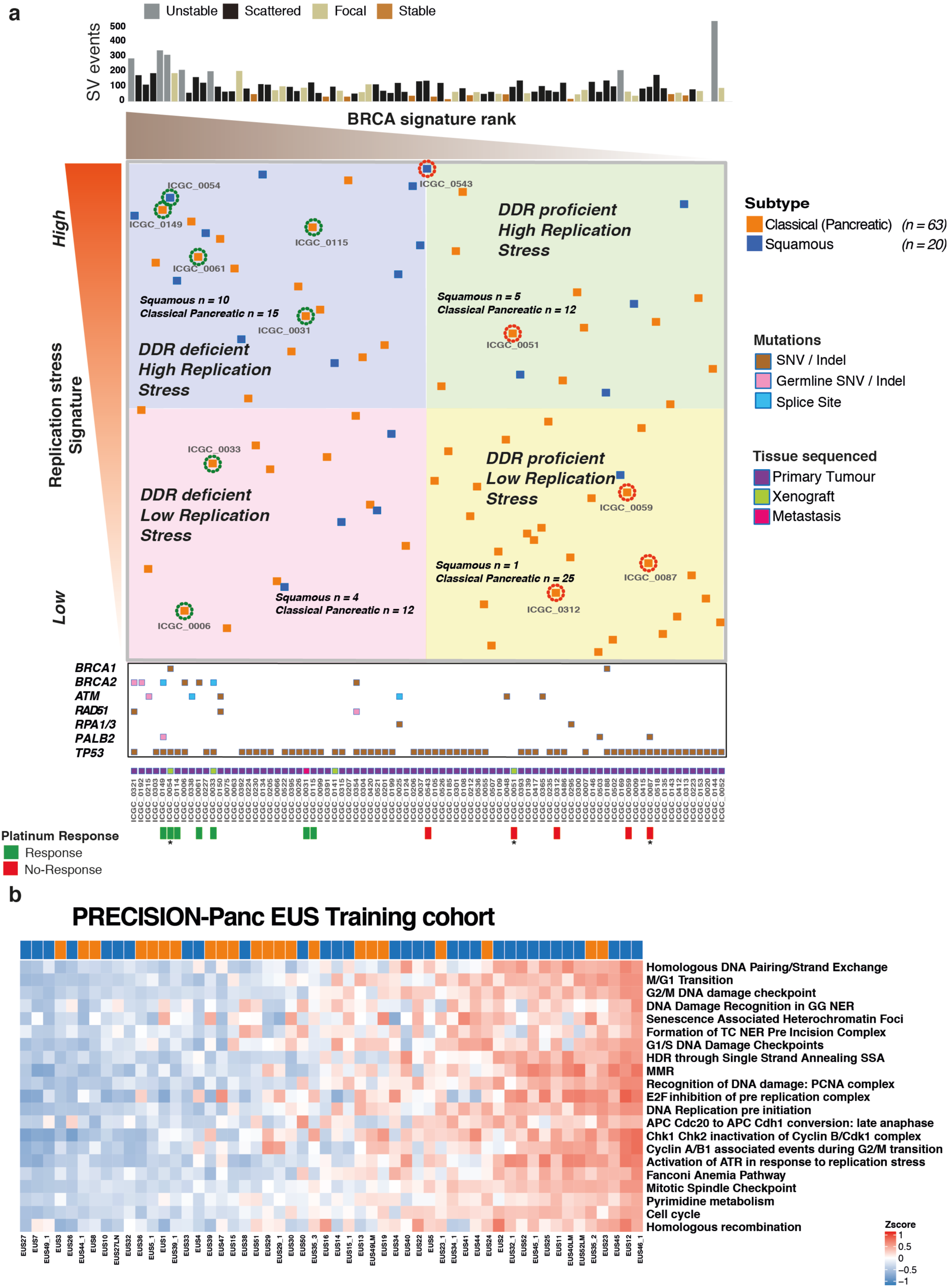
Targeting Replication Stress and DDR deficiency in clinical cohorts of PC. **a)** Bulk tumor samples from the ICGC PC cohort that have undergone both WGS and RNAseq are ranked from left to right based on the COSMIC BRCA mutational signature as a scale of DDR deficiency (x-axis) and top to bottom by the novel transcriptomic signature of replication stress (y-axis). HR pathway gene mutations and source of tissue sequenced is marked along the x-axis. Platinum response is marked along x-axis and related patient encircled at individual points where green represents response and red indicates resistance. * indicates PDX response data. Relevant molecular subtype frequency (squamous versus classical pancreatic) is indicated for each quadrant, demonstrating that Squamous PC was associated with high ranking replication stress score (15 out of 41 versus 5 out of 42) (*P* = 0.009; Chi-square test). **b)** The replication stress signature in the Precision-Panc endoscopic ultrasound FNB training cohort, demonstrating its clinical utility in the advanced disease setting (34% of cohort was locally advanced and 37% metastatic). The top-ranking quartile of replication stress signature score as high demonstrated that 50% of squamous tumors were within this group, compared to only 21% of the Classical Pancreatic tumors (*P* = 0.027, Chi square test).

The replication stress signature was then applied to The Cancer Genome Atlas (TCGA)^9^ high epithelial cellularity set (ABSOLUTE purity ≥0.2), and the ICGC micro-array transcriptomic data sets (Supplementary Figure 5). Again, the top-ranking quartile of replication stress signature was significantly enriched with squamous subtype PC (TCGA set *P* = 0.009, micro-array set *P* = 0.037) (Supplementary Figure 5). We then examined the potential clinical utility of the replication stress signature in biopsy material acquired through the Precision-Panc endoscopic ultrasound fine needle biopsy (FNB) training cohort (*n* = 54), recruited and collected during the development of the Precision-Panc (Figure 6b)^43^. As in the other cohorts, this demonstrated enrichment of the squamous subtype with high replication stress (*P* = 0.027) and provides proof-of-principle clinical validity that the signature can be generated from FNB material and be utilized as a putative biomarker in the clinical setting.

## Discussion

Identifying responsive patient subgroups is crucial to therapeutic development and improving outcomes for PC. Genomic sequencing studies and the development of novel therapeutic agents has made DNA damage response mechanisms one of the most attractive therapeutic opportunities in PC^4,8^. Using surrogate markers of DDR deficiency (HRD signature, structural variation, the COSMIC BRCA mutational signature, mutations in HR pathway genes) we demonstrate that DDR deficient PC respond preferentially to both platinum and PARP inhibitors in PDCLs (*n* = 15) and long lasting complete and near complete responses in a DDR deficient PDX models with single agent PARP inhibition with Olaparib. This was as effective as Cisplatin monotherapy, or combination treatment using Cisplatin and Olaparib suggesting that in appropriately selected patients, PARP inhibitor monotherapy can potentially induce clinically relevant responses similar to platinum. This provides potential therapeutic options for patients with poor performance status, or after intolerance or acquired resistance to platinum has developed^30^. Predicting platinum response is more complex than using point mutations in DDR genes and the COSMIC BRCA mutational signature alone. Structural variation signatures, including > 200 SVs^4^ and the SV pattern HRD signature appear to be robust, but require testing in clinical settings.

We define a novel replication stress signature, which is associated with the squamous subtype in PDCLs and bulk tumors from multiple PC cohorts (*n* = 383 patients). Elevated replication stress, as defined by this signature, is associated with functional deficiencies in DNA replication, leading to a therapeutic vulnerability as demonstrated by cell viability assays and siRNA functional screen. This molecular feature is independent of DDR deficiency and platinum response and offers patients with ‘DNA replication defects’ alternative therapeutic options to standard of care platinum chemotherapy. Based on part of these data, a biomarker driven therapeutic hypothesis was generated for testing on a series of clinical trials as part of the Precision-Panc platform (**Visual Abstract)**. Tumors that are DDR deficient can be targeted with platinum-based therapy, or in the context of a patient with reduced performance status or in the 2^nd^ line, PARP inhibitors. Patients with high replication stress can be targeted with ATR or WEE1 inhibitors, which can be combined with PARP inhibitors or platinum if concurrent DDR deficiency exist (**Visual Abstract)**, or after platinum resistance develops.

In summary, we develop and perform pre-clinical testing on novel biomarkers of DDR deficiency and replication stress which have potential clinical utility. Well-designed precision oncology platforms, such as Precision-Panc (precisionpanc.org), will enable biomarker driven clinical testing and allow refinement of biomarkers predicting meaningful responses and potential translation into clinical practice.

## Materials and methods

Full methods and references can be found in the supplementary material

### Human research ethics approvals

Ethical approval was obtained for all human samples and data (Supplementary Material).

### Cell culture

Patient derived cell lines (PDCLs) were generated as previously described^4,44–46^. PDCLs were cultured in conditions specifically formulated for each individual line based on growth preferences and those resulting in cell lines that most closely resembled physiological cells from the initial tumor. Detailed culture media formulations are given in Supplementary Material Table 1. Cells were grown in a humidified environment with either 5% or 2% CO^2^ at 37° (Supplementary Material Table 2). All cell lines were profiled by short tandem repeat (STR) DNA profiling as unique (CellBankaustralia.com). Cell lines were tested routinely for mycoplasma contamination using MycoAlert PLUS Mycoplasma Detection Kit (Lonza, #LT07 – 318).

### Reverse Phase Protein Array

Samples were lysed in RIPA lysis buffer (50mM Tris-HCL at pH7.4, 150mM Sodium Chloride, 5mM EGTA, 0.1% SDS, 1% NP40, 1% Deoxycholate, supplemented protease and phosphatase inhibitor tablets; Roche Applied Science Cat. #: 05056489001 and 04906837001) for 30 min on ice and cleared by centrifugation at 14K for 15 min at 4°C. Protein concentration was determined using a Bradford assay (Sigma) and all samples were normalized to 2mg/ml. 4x SDS sample buffer (40% Glycerol, 8% SDS, 0.25M Tris-HCL, pH 6.8. 1/10^th^ vol/vol 2-mercaptoethanol) was added to each sample, followed by incubation at 80°C for 5 mins. Serial dilutions (1, 0.5, 0.25, 0.125) were then prepared by diluting samples in PBS. Samples were printed onto Avid nitrocellulose coated glass slides (Grace Biolabs) using an Aushon 2470 microarrayer (Aushon Biosystems), with 2 technical replicates per sample. Slides were processed as follows: 4 × 15 min washes with dH20, incubated with antigen retrieval agent (Reblot strong, Millipore) for 15 min, 3 × 5 min washes with PBST, incubated with superblock TBST (ThermoFisher Scientific) for 10 min, 3 × 5 min washes with TBST, incubated with primary antibodies (all 1:200) diluted in superblock TBST for 60 mins, 3 × 5 min washes with TBST, blocked with superblock TBST for 10 mins, 3 × 5 min washes with TBST, incubated with anti-rabbit dylight 800 secondary Ab (1:2000 in superblock TBST)(Cell Signalling Technologies) for 30 mins, 3 × 5 min washes with TBST, 1 × 5 min wash with dH20, slides spun at 2000rpm for 5mins and allowed to air dry in the dark. An additional slide was stained with FAST Green FCF for normalization against total protein: 3 × 5min washes with dH20, incubated for 15 mins in 1% NaOH, slides rinsed 20 × in dH20, incubated for 10 min in dH20, incubated in de-stain (30% methanol, 7% glacial acetic acid, 63% dH20) for 15 min, incubated for 3 mins in FAST green staining solution (0.0025%w/v FAST green in de-stain), rinsed 20 × in dH20, incubated for 15 mins in de-stain solution, rinsed 20 x in dH20, spun at 2000rpm for 5 mins and allowed to air dry in the dark. All steps were performed at room temperature with agitation. Slides were visualized using an Innopsys 710AL infra-red microarray scanner and signals quantified using MAPIX microarray image analysis software (Innopsys). Non-specific signals were determined by omitting the primary antibody incubation step. All signals were within the linear range of detection with an R^2^ vlaue >0.9. Final output is the median value for each dilution series, background subtracted and normalized for protein loading.

### In Vitro Cytotoxicity assays

Cells were seeded on 96 well plates (Costar®, Corning Incorporated) and allowed to adhere for 24 hours. Cells were treated with increasing doses of Cisplatin (Accord Healthcare), AZD6738 (AstraZeneca®), AZD1775 (AstraZeneca®) and AZD7762 (AstraZeneca®) for 72hours. Cells were treated with BMN-673 (Pfizer Inc.), Rucaparib (Clovis Oncology), CFI-400945 (Cayman Chemical) and Palbociclib (Pfizer Inc.) for a total of 9 days, with repeated dosing every 72 hours in conjunction with changing cell media. Actinomycin D (Sigma), drug vehicle (DMSO) and media only controls were performed on each individual plate. For all other cytotoxicity assays, cells were plated in 96-well plates and treated with serial dilutions of indicated inhibitors 24hrs after plating for indicated time points. Cell viability was determined using CellTiter 96® Aqueous non-radioactive cell proliferation assay composed of solutions of a tetrazolium compound [3-(4,5-dimethylthiazol-2-yl)-5-(3-carboxymethoxyphenyl)-2-(4-sulfophenyl)-2H-tetrazolium, inner salt; MTS] and an electron-coupling reagent (phenazine methosulfate; PMS) (Promega, Madison, WI, USA). The assay was performed at an absorbance of 490 nm using an ELISA plate reader (Tecan Trading AG). Background absorbance was corrected for by wells containing medium alone and the absorbance was normalised to a scale of 0% (complete cell death by actinomycin D (5 - 10μg/ml) to 100% (no drug). At least 3 biological repeats were performed for each experiment. IC50 calculation and dose response curves were generated using GraphPad Prism 6 (GraphPad Software Inc, La Jolla CA).

### PDX

Patient derived xenografts (PDX) of PDAC were generated and comprehensively characterised as part of the ICGC project. BALB/c nude mice were anaesthetised and a single PDX fragment was inserted sub-cutaneously into the right flank according to standard operating procedure. PDX models were grown to 150mm^3^ (volume = length^2^ x width / 2), at this point each PDX was randomised to a different treatment regime. Responsive PDXs were treated once tumour size returned to 150mm^3^, up to a maximum of three rounds. Resistant models were treated after a treatment break of 2 weeks in accordance with current clinical treatment regimes, up to a maximum of 2 rounds. Each experiment was terminated once tumour volume reached end-point (750mm^3^), in accordance with home office animal welfare regulations. Full methods can be found in supplementary material.

### γH2AX and pRPA foci formation assay

PDCLs were cultured as standard and seeded in 96 well plates at a concentration of 10^4^ cells per well. At 24 hours after seeding, cells were either left untreated or exposed to 4 Gray (Gy) ionizing radiation (IR) and processed for analysis at 2, 4 and 20 hours after exposure. Cells were stained with primary antibodies at a dilution of 1:1000 with anti-pRPA32 (S4/S8, Bethyl Laboratories Inc.) and anti-γH2AX (Ser139, MERCK). Secondary antibodies used were Alexa 488 anti-mouse IgG (green) and Cy3 anti-rabbit IgG (Sigma). DAPI (Life Technologies) was used as a nuclear stain. Confocal imaging was performed using the Opera Phenix™ high content screening system (PerkinElmer) at 63x magnification using a water objective, at wavelengths of 405nm (DAPI), 488nm (Alexa) and 561nm (Cy3). A minimum of 320 cells (median 980, range 322 - 1886), in two separate experiments, were analyzed for each time point. Image analysis was performed using Columbus™ Image data storage and analysis system (PerkinElmer Inc, Waltham, MA). Statistical analysis was performed using GraphPad Prism 6 (GraphPad Software Inc, CA USA).

### Nucleic acid extraction

DNA and RNA extraction were performed using previously published methods^4^.

### Whole-genome library preparation

Whole-genome libraries were generated using either the Illumina TruSeq DNA LT sample preparation kit (Illumina, Part no. FC-121–2001 and FC-121–2001) or the Illumina TruSeq DNA PCR-free LT sample preparation kit (Illumina, Part no. FC-121–3001 and FC-121–3002) according to the manufacturer’s protocols with some modifications (Illumina, Part no. 15026486 Rev. C July 2012 and 15036187 Rev. A January 2013 for the two different kits respectively). Full methods can be found in the supplementary material.

### RNA sequencing library generation and sequencing

RNA-seq libraries were generated using TruSeq Stranded Total RNA (part no. 15031048 Rev. D April 2013) kits, using on a Perkin Elmer’s Sciclone G3 NGS Workstation (product no. SG3-31020-0300). Full methods can be found in the supplementary material.

### Library sequencing

All libraries were sequenced using the Illumina HiSeq 2000/2500 system with TruSeq SBS Kit v3 - HS (200-cycles) reagents (Illumina, Part no. FC-401-3001), to generate paired-end 101 bp reads.

### Copy number analysis

Matched tumour and normal patient DNA was assayed using Illumina SNP BeadChips as per manufacturer’s instructions (Illumina, San Diego CA) (HumanOmni1-Quad or HumanOmni2.5–8 BeadChips) and analysed as previously described.

### Identification and verification of structural variants

The Somatic structural variant pipeline were identified using the qSV tool. A detailed description of its use has been recently published^4,47^.

### Identification of and verification of point mutations

Substitutions and indels were called using a consensus calling approach that included qSNP, GATK and Pindel. The details of call integration and filtering, and verification using orthogonal sequencing and matched sample approaches are as previously described^4,47,48^.

### Mutational signatures

Mutational signatures were defined for genome-wide somatic substitutions, as previously described^4^.

### siRNA screening

Prior to siRNA screening, optimal cell number per well and optimal reverse transfection reagents for each PDCL were identified by assessing transfection efficiency, using six different transfection reagents (Dharmafect 1-4, RNAimax, Lipofectamine 2000), using the manufacturers’ instructions. Experimental conditions were selected that met the following criteria: (i) compared to a mock control (no lipid, no siRNA), the transfection of non-silencing negative control siRNA caused no more than 20 % cell inhibition; (ii) compared to non-silencing negative control siRNA, the transfection of PLK1–targeting siRNA caused more than 80% cell inhibition; (iii) cell confluency reached 70% within the range of 4-7 days. The later criteria allowed assays to be terminated whilst cells were in growth phase. Once optimal conditions were established, each PDCL was reverse transfected in a 384 well-plate format with a custom siGENOME siRNA library (Dharmacon, USA) designed to target 714 kinase coding genes, 256 protein phosphatase coding genes, 722 genes implicated in energy metabolism, 73 tumor suppressor genes and 166 genes involved in the repair of DNA damage (Supplementary Table 19 for list of genes covered in the siRNA library). Each well in the 384 well-plate arrayed library contained a SMARTpool of four distinct siRNA species targeting different sequences of the target transcript. Each plate was supplemented with non-targeting siCONTROL and siPLK1 siRNAs (Dharmacon, USA). Cell viability was estimated five days after transfection using a luminescent assay detecting cellular ATP levels (CellTitre-Glo, Promega). Luminescence values were processed using the cellHTS2 R package. To evaluate the effect of each siRNA pool on cell viability, we log2 transformed the luminescence measurements and then centred these to the median value for each plate. The plate-centred data were scaled to the median absolute deviation (MAD) of the library as a whole to produce robust Z-scores. All screens were performed in triplicate. Screens judged to have poor dynamic range (Z’ factor < 0) or poorly correlated replicates (r < 0.7) were excluded during an evaluation of screen quality. Z scores were adjusted using a quantile normalization.

### siRNA screen analysis

siRNA “hits” were identified by calculating the median absolute deviation of normalized Z-scores for a given siRNA across all samples and identifying sample Z scores greater than or equal to 2 x the median absolute deviation. This analysis generated a “seed” matrix (n siRNA hits x m samples) which was used as starting input for the Randon Walk with Restart (RWR) algorithm as implemented by the R package *dnet*. This algorithm was used to identify functionally important subnetworks associated with cell viability from a curated protein-protein interaction network STRING v 10. Considering the complex nature of topological features of human interactome data, we introduce a randomization-based test to evaluate the candidate interactors utilizing 1000 topologically matched random networks. Candidate interactors that remain significant (i.e., *p* _edge_<0.05) were identified and a consensus subnetwork was constructed by collapsing all sample-specific results. The resulting network was visualized using RedeR.

### RNAseq analysis

RNA-seq read mapping was performed using the bcbio-nextgen project RNAseq pipeline (https://bcbio-nextgen.readthedocs.org/en/latest/). Briefly, after quality control and adaptor trimming, reads were aligned to the GRCh37 genome build using STAR Counts for known genes were generated using the function featureCounts in the R/Bioconductor package “Rsubread”. The R/Bioconductor package “DESeq2” was used to normalize count data between samples and to identify differentially expressed genes. Expression data were normalized using the rlog transform in the DESeq2 package and these values were used for all downstream analyses.

### WGCNA analysis

Weighted gene co-expression network analysis (WGCNA) was used to generate a transcriptional network from rlog normalized RNAseq data. Briefly, WGCNA clusters genes into network modules using a topological overlap measure (TOM). The TOM is a highly robust measure of network interconnectedness and essentially provides a measure of the connection strength between two adjacent genes and all other genes in a network. Genes are clustered using 1-TOM as the distance measure and gene modules are defined as branches of the resulting cluster tree using a dynamic branch-cutting algorithm. Full methods can be found in the supplementary material.

### Identification of significant subtype specific changes in pathways and/or processes

The R package *clipper* was used to identify pathways and/or processes showing significant change between PDCL subtypes. *clipper* implements a two-step empirical approach, employing a statistical analysis of means and concentration matrices of graphs derived from pathway topologies, to identify signal paths having the greatest association with a specific phenotype.

### Pathway analysis

Ontology and pathway enrichment analysis was performed using the R package ‘dnet’ and/or the ClueGO/CluePedia Cytoscape plugins as indicated. Visualisation and/or generation of network diagrammes was performed using either Cytoscape or the R package *RedeR*.

### Homologous Recombination Signature generation

Signature generation was done using WGS data as previously described^31,32^. A positive signature was defined as if samples had > 50 SVs and % deletions, duplication or translocations > 70, and structural variation pattern was not focal (ie large number of SVs not due to chromothripsis or amplifications). Plus, either the predominant variant types were deletions and translocations and the median deletion size was < 10kb OR the predominant variant type was duplication and the median duplication size was < 50 kb.

### Replication Stress signature generation

Differentially expressed genes were compared to genes associated with gene ontology (GO) terms using the R package ‘dnet’. Significantly expressed GO terms involved in DNA damage response and cell cycle control were selected. Differential expression of each selected GO term was applied to each individual PDCL that underwent RNA sequencing. This in turn, was used to generate a composite score by totaling the score for each selected GO term. The ‘*sig.score*’ function from the R package *genefu* was used to calculate a specific signature score in a given sample using the signatures generated for each pathway and/or process. This was termed the replication stress signature. Generation of signature score for bulk tumor samples followed the same methodology

### Bulk expression sets and immune signature scores

Bulk RNAseq expression data, subtype assignments and immune signature scores were obtained from Bailey *et al*^5^.

### Gene set enrichment of PDAC subtypes

Gene set enrichment was performed using the R package ‘GSVA’. Gene sets representing PDAC subtypes were generated as previously described.

### Clustering and subtype assignment

The package ‘ConsensusClusterPlus’ was used to classify PDCLs according to the expression signatures defined in Moffitt *et al.*^18^ and Bailey *et al.*^5^ Gene sets representing PDAC subtypes were generated as previously described.

### Statistical analysis

A Kruskal–Wallis test was applied to the indicated stratified scores to determine whether distributions were significantly different. Fisher’s exact tests were used to evaluate the association between dichotomous variables.

### Plot generation

Heatmaps and oncoplots were generated using the R package *ComplexHeatmap.* Dotcharts, density plots and boxplots were generated using the R package *ggpubr.* Violin plots were generated using the python package *Seaborn.* Biplot was generated using the R package *ggfortify*. All other plots were generated using the R package *ggplot2*.

## Funding support

National Health and Medical Research Council of Australia (NHMRC; 631701, 535903, 427601)

Queensland Government (NIRAP)

University of Queensland Institute for Molecular Bioscience

Australian Government: Department of Innovation, Industry, Science and Research (DIISR)

Australian Cancer Research Foundation (ACRF)

Cancer Council NSW: (SRP06-01, SRP11-01. ICGC)

Cancer Institute NSW: (10/ECF/2-26; 06/ECF/1-24; 09/CDF/2-40; 07/CDF/1-03; 10/CRF/1-01, 08/RSA/1-15, 07/CDF/1-28, 10/CDF/2-26,10/FRL/2-03, 06/RSA/1-05, 09/RIG/1-02, 10/TPG/1-04, 11/REG/1-10, 11/CDF/3- 26)

Garvan Institute of Medical Research

Avner Nahmani Pancreatic Cancer Research Foundation

R.T. Hall Trust

Petre Foundation

Philip Hemstritch Foundation

Gastroenterological Society of Australia (GESA)

American Association for Cancer Research (AACR) Landon Foundation – INNOVATOR Award

Royal Australasian College of Surgeons (RACS)

Royal Australasian College of Physicians (RACP)

Royal College of Pathologists of Australasia (RCPA)

Susan Wojcicki and Dennis Tropper

NIH grant CA62924

NIH grant 5R01CA150190-07

NIH grant P50 CA102701

Cancer Research UK C29717/A17263, C29717/A18484, C596/A18076, C596/A20921, A23526

Wellcome Trust Senior Investigator Award: 103721/Z/14/Z

Pancreatic Cancer UK Future Research Leaders Fund FLF2015_04_Glasgow

Scottish Genome Partnership SEHHD-CSO 1175759/2158447

MRC/EPSRC Glasgow Molecular Pathology Node

The Howat Foundation

Italian Cancer Genome Project – Ministry of University [FIRB RBAP10AHJB];

Associazione Italiana Ricerca Cancro [grant number: 12182];

FP7 European Community Grant Cam-Pac [no: 602783];

Italian Ministry of Health [FIMP- CUP_J33G13000210001].

ICGC Ontario Institute for Cancer Research

## Visual Abstract

**Targeting DNA damage response and replication stress in PC: translating pre-clinical discovery to clinical trials in PC**

Whole genome and transcriptome data were utilized to generate novel, and test previously described, putative biomarkers of DDR deficiency and replication stress. This informed pre-clinical therapeutic sensitivity, which in turn informed biomarker hypotheses for clinical testing on the Precision-Panc platform.

## References

1 Siegel, R. L., Miller, K. D. & Jemal, A. Cancer statistics, 2019. CA: a cancer journal for clinicians 69, 7–34, doi:10.3322/caac.21551 (2019).

2 Rahib, L. et al. Projecting cancer incidence and deaths to 2030: the unexpected burden of thyroid, liver, and pancreas cancers in the United States. Cancer research 74, 2913–2921, doi:10.1158/0008-5472.CAN-14-0155 (2014).

3 Biankin, A. V. et al. Pancreatic cancer genomes reveal aberrations in axon guidance pathway genes. Nature 491, 399–405, doi:10.1038/nature11547 (2012).

4 Waddell, N. et al. Whole genomes redefine the mutational landscape of pancreatic cancer. Nature 518, 495–501, doi:10.1038/nature14169 (2015).

5 Bailey, P. et al. Genomic analyses identify molecular subtypes of pancreatic cancer. Nature 531, 47–52, doi:10.1038/nature16965 (2016).

6 Humphris, J. L. et al. Hypermutation In Pancreatic Cancer. Gastroenterology 152, 68–74 e62, doi:10.1053/j.gastro.2016.09.060 (2017).

7 Witkiewicz, A. K. et al. Whole-exome sequencing of pancreatic cancer defines genetic diversity and therapeutic targets. Nature communications 6, 6744, doi:10.1038/ncomms7744 (2015).

8 Dreyer, S. B., Chang, D. K., Bailey, P. & Biankin, A. V. Pancreatic Cancer Genomes: Implications for Clinical Management and Therapeutic Development. Clinical cancer research: an official journal of the American Association for Cancer Research 23, 1638–1646, doi:10.1158/1078-0432.CCR-16-2411 (2017).

9 Cancer Genome Atlas Network. Integrated Genomic Characterization of Pancreatic Ductal Adenocarcinoma. Cancer cell 32, 185–203 e113, doi:10.1016/j.ccell.2017.07.007 (2017).

10 Notta, F. et al. A renewed model of pancreatic cancer evolution based on genomic rearrangement patterns. Nature 538, 378–382, doi:10.1038/nature19823 (2016).

11 Jones, S. et al. Core signaling pathways in human pancreatic cancers revealed by global genomic analyses. Science (New York, N.Y.) 321, 1801–1806, doi:10.1126/science.1164368 (2008).

12 Biankin, A. V., Piantadosi, S. & Hollingsworth, S. J. Patient-centric trials for therapeutic development in precision oncology. Nature 526, 361–370, doi:10.1038/nature15819 (2015).

13 Biankin, A. V. & Maitra, A. Subtyping Pancreatic Cancer. Cancer cell 28, 411–413, doi:10.1016/j.ccell.2015.09.020 (2015).

14 Collisson, E. A. et al. Subtypes of pancreatic ductal adenocarcinoma and their differing responses to therapy. Nature medicine 17, 500–503, doi:10.1038/nm.2344 (2011).

15 Connor, A. A. et al. Association of Distinct Mutational Signatures With Correlates of Increased Immune Activity in Pancreatic Ductal Adenocarcinoma. JAMA Oncol 3, 774–783, doi:10.1001/jamaoncol.2016.3916 (2017).

16 Maurer, C. et al. Experimental microdissection enables functional harmonisation of pancreatic cancer subtypes. Gut, gutjnl-2018-317706, doi:10.1136/gutjnl-2018-317706 (2019).

17 Puleo, F. et al. Stratification of Pancreatic Ductal Adenocarcinomas Based on Tumor and Microenvironment Features. Gastroenterology, doi:10.1053/j.gastro.2018.08.033 (2018).

18 Moffitt, R. A. et al. Virtual microdissection identifies distinct tumor-and stroma-specific subtypes of pancreatic ductal adenocarcinoma. Nat Genet 47, 1168–1178, doi:10.1038/ng.3398 (2015).

19 Collisson, E. A., Bailey, P., Chang, D. K. & Biankin, A. V. Molecular subtypes of pancreatic cancer. Nature Reviews Gastroenterology & Hepatology, doi:10.1038/s41575-019-0109-y (2019).

20 Dreyer, S. B. et al. Precision Oncology in Surgery: Patient Selection for Operable Pancreatic Cancer. Annals of surgery, doi:10.1097/SLA.0000000000003143 (2018).

21 Zhang, J., Dai, Q., Park, D. & Deng, X. Targeting DNA Replication Stress for Cancer Therapy. Genes (Basel) 7, doi:10.3390/genes7080051 (2016).

22 Lecona, E. & Fernandez-Capetillo, O. Targeting ATR in cancer. Nature reviews. Cancer, doi:10.1038/s41568-018-0034-3 (2018).

23 Hanahan, D. & Weinberg, R. A. Hallmarks of cancer: the next generation. Cell 144, 646–674, doi:10.1016/j.cell.2011.02.013 (2011).

24 Di Micco, R. et al. Oncogene-induced senescence is a DNA damage response triggered by DNA hyper-replication. Nature 444, 638–642, doi:10.1038/nature05327 (2006).

25 Macheret, M. & Halazonetis, T. D. Intragenic origins due to short G1 phases underlie oncogene-induced DNA replication stress. Nature 555, 112–116, doi:10.1038/nature25507 (2018).

26 Conroy, T. et al. Unicancer GI PRODIGE 24/CCTG PA.6 trial: A multicenter international randomized phase III trial of adjuvant mFOLFIRINOX versus gemcitabine (gem) in patients with resected pancreatic ductal adenocarcinomas. Journal of Clinical Oncology 36, LBA4001–LBA4001, doi:10.1200/JCO.2018.36.18_suppl.LBA4001 (2018).

27 Murphy, J. E. et al. Total Neoadjuvant Therapy With FOLFIRINOX Followed by Individualized Chemoradiotherapy for Borderline Resectable Pancreatic Adenocarcinoma: A Phase 2 Clinical Trial. JAMA Oncol 4, 963–969, doi:10.1001/jamaoncol.2018.0329 (2018).

28 Conroy, T. et al. FOLFIRINOX versus gemcitabine for metastatic pancreatic cancer. The New England journal of medicine 364, 1817–1825, doi:10.1056/NEJMoa1011923 (2011).

29 Shroff, R. T. et al. Rucaparib Monotherapy in Patients With Pancreatic Cancer and a Known Deleterious BRCA Mutation. JCO Precis Oncol 2018, doi:10.1200/PO.17.00316 (2018).

30 Golan, T. et al. Maintenance Olaparib for Germline BRCA-Mutated Metastatic Pancreatic Cancer. The New England journal of medicine, doi:10.1056/NEJMoa1903387 (2019).

31 McBride, D. J. et al. Tandem duplication of chromosomal segments is common in ovarian and breast cancer genomes. The Journal of pathology 227, 446–455, doi:10.1002/path.4042 (2012).

32 Ng, C. K. et al. The role of tandem duplicator phenotype in tumour evolution in high-grade serous ovarian cancer. The Journal of pathology 226, 703–712, doi:10.1002/path.3980 (2012).

33 Kotsantis, P., Petermann, E. & Boulton, S. J. Mechanisms of Oncogene-Induced Replication Stress: Jigsaw Falling into Place. Cancer discovery 8, 537–555, doi:10.1158/2159-8290.CD-17-1461 (2018).

34 Karnitz, L. M. & Zou, L. Molecular Pathways: Targeting ATR in Cancer Therapy. Clinical cancer research: an official journal of the American Association for Cancer Research 21, 4780–4785, doi:10.1158/1078-0432.ccr-15-0479 (2015).

35 Zhang, Y. et al. Targeting radioresistant breast cancer cells by single agent CHK1 inhibitor via enhancing replication stress. Oncotarget 7, 34688–34702, doi:10.18632/oncotarget.9156 (2016).

36 Zheng, H., Shao, F., Martin, S., Xu, X. & Deng, C. X. WEE1 inhibition targets cell cycle checkpoints for triple negative breast cancers to overcome cisplatin resistance. Scientific reports 7, 43517, doi:10.1038/srep43517 (2017).

37 Yap, T. A. et al. Phase I modular study of AZD6738, a novel oral, potent and selective ataxia telangiectasia Rad3-related (ATR) inhibitor in combination (combo) with carboplatin, olaparib or durvalumab in patients (pts) with advanced cancers. European Journal of Cancer 69, S2, doi:10.1016/S0959-8049(16)32607-7 (2018).

38 Do, K. et al. Phase I Study of Single-Agent AZD1775 (MK-1775), a Wee1 Kinase Inhibitor, in Patients With Refractory Solid Tumors. Journal of clinical oncology: official journal of the American Society of Clinical Oncology 33, 3409–3415, doi:10.1200/JCO.2014.60.4009 (2015).

39 Dillon, M. T. et al. PATRIOT: A phase I study to assess the tolerability, safety and biological effects of a specific ataxia telangiectasia and Rad3-related (ATR) inhibitor (AZD6738) as a single agent and in combination with palliative radiation therapy in patients with solid tumours. Clin Transl Radiat Oncol 12, 16–20, doi:10.1016/j.ctro.2018.06.001 (2018).

40 Krebs, M. et al. 413PDPhase I clinical and translational evaluation of AZD6738 in combination with durvalumab in patients (pts) with lung or head and neck carcinoma. Annals of Oncology 29, doi:10.1093/annonc/mdy279.401 (2018).

41 Leijen, S. et al. Phase II Study of WEE1 Inhibitor AZD1775 Plus Carboplatin in Patients With TP53-Mutated Ovarian Cancer Refractory or Resistant to First-Line Therapy Within 3 Months. Journal of clinical oncology: official journal of the American Society of Clinical Oncology 34, 4354–4361, doi:10.1200/JCO.2016.67.5942 (2016).

42 Leijen, S. et al. Phase I Study Evaluating WEE1 Inhibitor AZD1775 As Monotherapy and in Combination With Gemcitabine, Cisplatin, or Carboplatin in Patients With Advanced Solid Tumors. Journal of clinical oncology: official journal of the American Society of Clinical Oncology 34, 4371–4380, doi:10.1200/JCO.2016.67.5991 (2016).

43 Dreyer, S. B. et al. Feasibility and clinical utility of endoscopic ultrasound guided biopsy of pancreatic cancer for next-generation molecular profiling. Chin Clin Oncol 8, 16, doi:10.21037/cco.2019.04.06 (2019).

44 Ruckert, F. et al. Five primary human pancreatic adenocarcinoma cell lines established by the outgrowth method. The Journal of surgical research 172, 29–39, doi:10.1016/j.jss.2011.04.021 (2012).

45 Chou, A. et al. Tailored first-line and second-line CDK4-targeting treatment combinations in mouse models of pancreatic cancer. Gut, doi:10.1136/gutjnl-2017-315144 (2017).

46 Pal, K. et al. Inhibition of endoglin-GIPC interaction inhibits pancreatic cancer cell growth. Molecular cancer therapeutics 13, 2264–2275, doi:10.1158/1535-7163.Mct-14-0291 (2014).

47 Nones, K. et al. Genomic catastrophes frequently arise in esophageal adenocarcinoma and drive tumorigenesis. Nature communications 5, 5224, doi:10.1038/ncomms6224 (2014).

48 Patch, A. M. et al. Whole-genome characterization of chemoresistant ovarian cancer. Nature 521, 489–494, doi:10.1038/nature14410 (2015).

